# UNCROSS: Filtering of high-frequency cross-talk in 16S amplicon reads

**DOI:** 10.1101/088666

**Authors:** Robert C. Edgar

## Abstract

Next-generation amplicon sequencing is widely used for surveying biological diversity in applications such as microbial metagenomics, immune system repertoire analysis and targeted tumor sequencing of cancer-associated genes. In such studies, assignment of reads to incorrect samples (cross-talk) is a well-documented problem that is rarely considered in practice. By considering unexpected OTUs in artificial (mock) samples, I estimate that cross-talk occurred for ~2% of the reads in one Illumina GAIIx run and eleven Illumina MiSeq runs targeting 16S ribosomal RNA. I also describe UNCROSS, an algorithm for detecting and filtering cross-talk in OTU tables.

## Introduction

Recent examples of next-generation amplicon sequencing experiments include the Human Microbiome Project (HMP Consortium, 2012), an analysis of the response of the human immune system to influenza vaccination (Jiang *et al.*, 2013) and a high-throughput search for known cancer-relevant variants in 16 oncogenes (Hadd *et al.*, 2013). In such studies, samples are multiplexed into a single run by embedding index sequences into amplicons to identify the sample of origin. Index sequences are sometimes called tags or barcodes, but I will avoid the latter terms here as some authors use them to refer to the biological sequence in an amplicon. An index sequence can be annealed to the start of the amplicon (Caporaso *et al.*, 2011; Derakhshani *et al.*, 2016) (*single-indexing*), while *dual-index* schemes attach indexes to both the ends of the construct (Kozich *et al.*, 2013; Derakhshani *et al.*, 2016). Previous studies have revealed unexpectedly high rates of cross-talk in both 454 (Carlsen *et al.*, 2012) and Illumina (Kircher *et al.*, 2012; Nelson *et al.*, 2014) data. Indexing methods designed to mitigate cross-talk have recently been proposed by (Esling *et al.*, 2015) and (Schnell *et al.*, 2015). Here, I investigate cross-talk in reads from one Illumina GAIIx run (Caporaso *et al.*, 2011) and eleven MiSeq paired-end sequencing runs (Kozich *et al.*, 2013) targeting the 16S gene. I describe UNCROSS, an algorithm for cross-talk detection in OTU tables, and show that it successfully identifies ~80% of spurious OTU entries due to cross-talk in these runs.

## Results

GAIIx reads were kindly provided by the authors as they are not deposited in the Short Read Archive as stated in Caporaso *et al*. 2011. They include 25 *in vivo* samples from different environments and three replicates of a designed (*mock*) community containing 67 strains. A single-index scheme was used with a 6-base index sequence. I created a partial reference database of 16S sequences for the mock community by matching species names to the Living Tree Project subset of the SILVA database (Yilmaz *et al.*, 2014). These sequences may have some differences compared to strains in the mock samples. I was unable to find reference sequences for nine of the species in the community.

MiSeq reads were obtained from <u>http://www.mothur.org/MiSeqDevelopmentData.html</u>, accessed 10th Jan. 2016. These are paired-end reads from eleven different MiSeq runs using three different versions of the Illumina Real-Time Analysis (RTA) and MiSeq Control Software (MCS) (Table S1 in Kozich *et al*., 2013). Twelve samples were sequenced in each run: three replicates of a mock sample with 21 species which was designed (Haas *et al.*, 2011) for the Human Microbiome Project, plus three replicates obtained from human gut, mouse gut and soil samples, respectively. A reference database for the HMP mock community was included in the download. A dual-index scheme was used allowing up to 96 distinct samples.

I created OTUs using UPARSE (Edgar, 2013). MiSeq read pairs were merged using a Bayesian assembler to ensure that consensus base calls and quality scores are correctly calculated in the overlapping segment (Edgar and Flyvbjerg, 2014). Quality filtering was performed using a maximum expected error threshold of one so that the most probable number of errors in each merged read is zero according to its quality scores (Edgar and Flyvbjerg, 2014). Merged reads with lengths <230nt or >270nt were discarded to select the V4 hypervariable region. An OTU table was generated by aligning reads to OTU sequences using USEARCH (Edgar, 2010). A read was assigned to the OTU with highest identity, or discarded if the top hit had <97% identity. For GAIIx reads, sample names were obtained by requiring an identical match to an index sequence. The posted MiSeq reads were already demultiplexed so sample identifiers were taken from the FASTQ filenames; for example, reads in Soil3_S6_L001_R1_001.fastq were assigned to sample Soil3. OTUs were classified by comparing their sequences first to the mock community reference database, then to SILVA (Pruesse *et al.*, 2007) if a match with ³97% identity was not found.

In all datasets, mock samples were found to have many more OTUs than expected from the designed community composition. In the GAIIx reads, 1,522 OTUs have one or more reads assigned to the mock samples, far more than the ~45 clusters obtained by clustering the known V4 sequences at 97% identity. In the MiSeq data, the runs have up to 727 mock OTUs with nine of the eleven datasets having >200 (Table 3). In all twelve datasets, most of the unexpected mock OTUs (i.e., those which do not match a reference sequence for a designed strain) have high abundances in the environmental samples and most or all these are therefore probably due to cross-talk.

Table 1 shows the 25 OTUs from run 130417 with the highest mock abundances. The unexpected mock OTUs (i.e., those which do not match a designed strain) have high abundance elsewhere. For example, OTU EF400979 has 73,265 reads in human gut and 393 reads in the mock samples. Similarly, all of the OTUs with high abundance in the mock samples are often found in low abundance in the environmental samples which is also strongly suggestive of cross-talk, though this is less clear as several of the mock species are human pathogens and thus could plausibly be present *in vivo*, especially in human gut. In Table 1, OTU table entries were annotated manually as *cross-talk*, *valid*, *contaminant* or *overlaid* by considering the most likely explanations for the reported counts. An *overlaid* entry is inferred to be present in both mock and environmental samples.

**Table 1.**
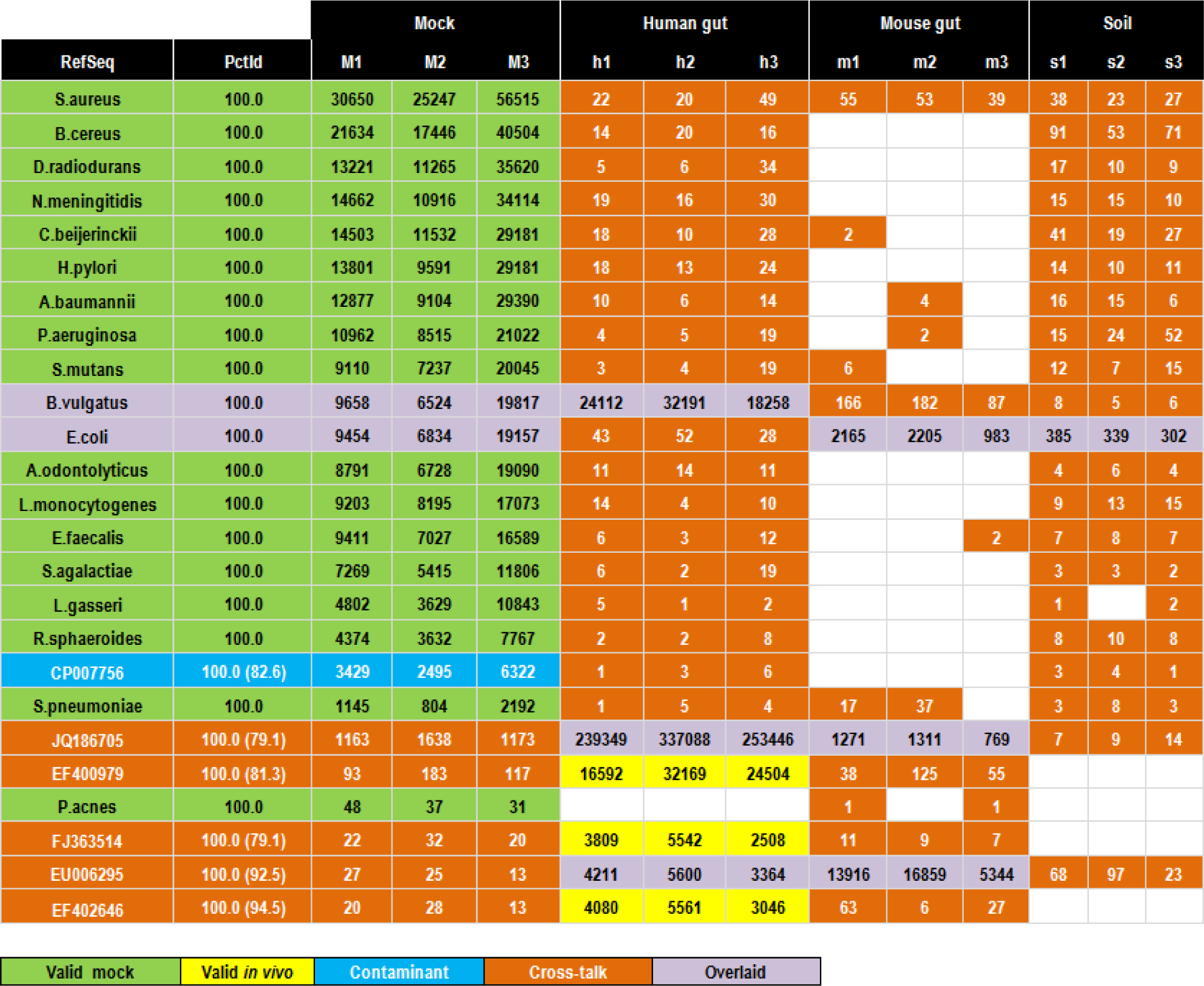
The 25 OTUs from MiSeq run 130417 with highest mock abundances. OTUs are sorted in order of decreasing total mock abundance. Counts are manually annotated as *valid* (green for mock, yellow for environmental), *contaminant* (blue, can be detected in mock only), *cross-talk* (orange) or *overlaid* (purple, meaning that the OTU is valid in the mock samples and one or more environmental sample). Reference sequences with species names (green) are designed strains in the mock community, otherwise are Genbank identifiers (blue for contaminant, purple for overlaid and orange for cross-talk). The PctId column gives identity with the reference sequence, values in parenthesis are identities with the most similar mock reference sequence to confirm that the contaminant and cross-talk counts are not due to noisy reads of expected strains. Note that two cross-talk OTUs (JQ186705 and EF400979) and one contaminant OTU (CP007756) have higher abundance than the least abundant expected strain (*P. acnes*).

| RefSeq | PctId | Mock |  |  | Human gut |  |  | Mouse gut |  |  | Soil |  |  |
| --- | --- | --- | --- | --- | --- | --- | --- | --- | --- | --- | --- | --- | --- |
|  |  | M1 | M2 | M3 | h1 | h2 | h3 | m1 | m2 | m3 | s1 | s2 | s3 |
| *S.aureus* | 100.0 | 30650 | 25247 | 56515 | 22 | 20 | 49 | 55 | 53 | 39 | 38 | 23 | 27 |
| *B.cereus* | 100.0 | 21634 | 17446 | 40504 | 14 | 20 | 16 |  |  |  | 91 | 53 | 71 |
| *D.radiodurans* | 100.0 | 13221 | 11265 | 35620 | 5 | 6 | 34 |  |  |  | 17 | 10 | 9 |
| *N.meningitidis* | 100.0 | 14662 | 10916 | 34114 | 19 | 16 | 30 |  |  |  | 15 | 15 | 10 |
| *C.beijerinckii* | 100.0 | 14503 | 11532 | 29181 | 18 | 10 | 28 | 2 |  |  | 41 | 19 | 27 |
| *H.pylori* | 100.0 | 13801 | 9591 | 29181 | 18 | 13 | 24 |  |  |  | 14 | 10 | 11 |
| *A.baumannii* | 100.0 | 12877 | 9104 | 29390 | 10 | 6 | 14 |  | 4 |  | 16 | 15 | 6 |
| *P.aeruginosa* | 100.0 | 10962 | 8515 | 21022 | 4 | 5 | 19 |  | 2 |  | 15 | 24 | 52 |
| *S.mutans* | 100.0 | 9110 | 7237 | 20045 | 3 | 4 | 19 | 6 |  |  | 12 | 7 | 15 |
| *B.vulgatus* | 100.0 | 9658 | 6524 | 19817 | 24112 | 32191 | 18258 | 166 | 182 | 87 | 8 | 5 | 6 |
| *E.coli* | 100.0 | 9454 | 6834 | 19157 | 43 | 52 | 28 | 2165 | 2205 | 983 | 385 | 339 | 302 |
| *A.odontolyticus* | 100.0 | 8791 | 6728 | 19090 | 11 | 14 | 11 |  |  |  | 4 | 6 | 4 |
| *L.monocytogenes* | 100.0 | 9203 | 8195 | 17073 | 14 | 4 | 10 |  |  |  | 9 | 13 | 15 |
| *E.faecalis* | 100.0 | 9411 | 7027 | 16589 | 6 | 3 | 12 |  |  | 2 | 7 | 8 | 7 |
| *S.agalactiae* | 100.0 | 7269 | 5415 | 11806 | 6 | 2 | 19 |  |  |  | 3 | 3 | 2 |
| *L.gasseri* | 100.0 | 4802 | 3629 | 10843 | 5 | 1 | 2 |  |  |  | 1 |  | 2 |
| *R.sphaeroides* | 100.0 | 4374 | 3632 | 7767 | 2 | 2 | 8 |  |  |  | 8 | 10 | 8 |
| CP007756 | 100.0 (82.6) | 3429 | 2495 | 6322 | 1 | 3 | 6 |  |  |  | 3 | 4 | 1 |
| *S.pneumoniae* | 100.0 | 1145 | 804 | 2192 | 1 | 5 | 4 | 17 | 37 |  | 3 | 8 | 3 |
| JQ186705 | 100.0 (79.1) | 1163 | 1638 | 1173 | 239349 | 337088 | 253446 | 1271 | 1311 | 769 | 7 | 9 | 14 |
| EF400979 | 100.0 (81.3) | 93 | 183 | 117 | 16592 | 32169 | 24504 | 38 | 125 | 55 |  |  |  |
| *P.acnes* | 100.0 | 48 | 37 | 31 |  |  |  | 1 |  | 1 |  |  |  |
| FJ363514 | 100.0 (79.1) | 22 | 32 | 20 | 3809 | 5542 | 2508 | 11 | 9 | 7 |  |  |  |
| EU006295 | 100.0 (92.5) | 27 | 25 | 13 | 4211 | 5600 | 3364 | 13916 | 16859 | 5344 | 68 | 97 | 23 |
| EF402646 | 100.0 (94.5) | 20 | 28 | 13 | 4080 | 5561 | 3046 | 63 | 6 | 27 |  |  |  |

|  |  |  |  |  |
| --- | --- | --- | --- | --- |
| Valid mock | Valid *in vivo* | Contaminant | Cross-talk | Overlaid |

Table 2 shows manual annotations for the 25 most abundant OTUs in the GAIIx data. Notably, none of the 400 counts in this table are zero despite the different environments and the fact that most OTUs are not expected in the mock samples. This can be explained by observing that a large majority (361/400 = 90%) of the OTU table entries are consistent with cross-talk and most of these should therefore probably be zero. The correct number of non-zero counts in these 25 OTUs is estimated to be approximately 400–361 = 39, an order of magnitude fewer than the 400 obtained without correcting for cross-talk.

**Table 2.**
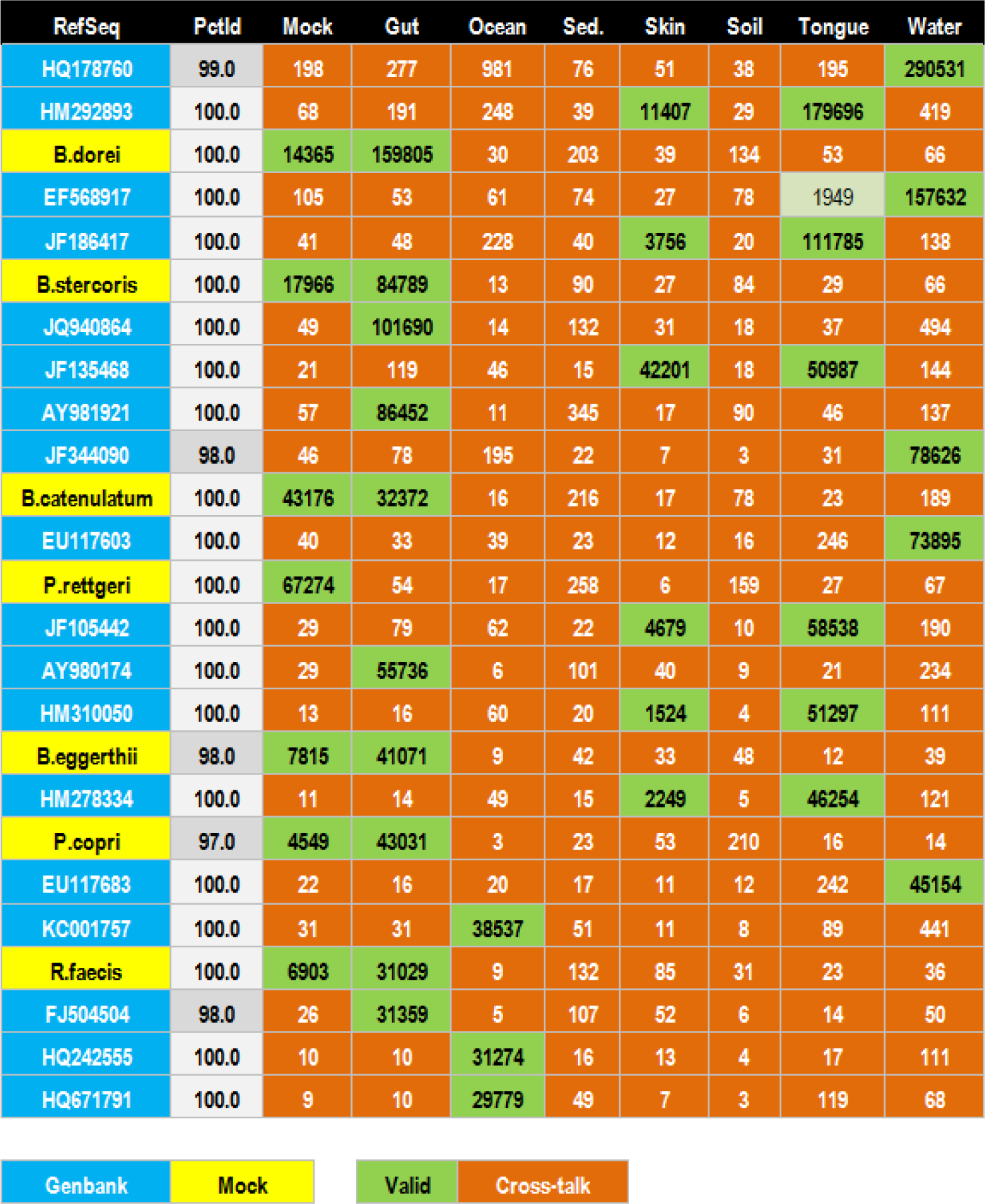
The top 25 most abundant OTUs from the GAIIx run. Most entries in this table are probably spurious due to cross-talk and should therefore be zero. OTUs are sorted in order of decreasing total abundance. Counts are manually annotated as *valid* (green) or *cross-talk* (orange). Most cases are readily classified except Tongue in the fourth OTU (light green) which has 1949/159979=1.2% of the total reads, which could be cross-talk but is a distinctly higher fraction than other probable cross-talk counts seen in the table. Reference sequences with species names (yellow) are designed strains in the mock community, otherwise are Genbank identifiers (blue). Nine species are missing from the mock reference database, so some, but not all, of the OTUs marked with Genbank identifiers may be expected mock OTUs. PctId gives the OTU identity with the reference sequences. Two of the mock identities are <100% which is probably due to reference sequences which do not match exactly because they were obtained from different strains of the same species.

| RefSeq | PctId | Mock | Gut | Ocean | Sed. | Skin | Soil | Tongue | Water |
| --- | --- | --- | --- | --- | --- | --- | --- | --- | --- |
| HQ178760 | 99.0 | 198 | 277 | 981 | 76 | 51 | 38 | 195 | 290531 |
| HM292893 | 100.0 | 68 | 191 | 248 | 39 | 11407 | 29 | 179696 | 419 |
| **B.dorei** | 100.0 | 14365 | 159805 | 30 | 203 | 39 | 134 | 53 | 66 |
| EF568917 | 100.0 | 105 | 53 | 61 | 74 | 27 | 78 | 1949 | 157632 |
| JF186417 | 100.0 | 41 | 48 | 228 | 40 | 3756 | 20 | 111785 | 138 |
| **B.stercoris** | 100.0 | 17966 | 84789 | 13 | 90 | 27 | 84 | 29 | 66 |
| JQ940864 | 100.0 | 49 | 101690 | 14 | 132 | 31 | 18 | 37 | 494 |
| JF135468 | 100.0 | 21 | 119 | 46 | 15 | 42201 | 18 | 50987 | 144 |
| AY981921 | 100.0 | 57 | 86452 | 11 | 345 | 17 | 90 | 46 | 137 |
| JF344090 | 98.0 | 46 | 78 | 195 | 22 | 7 | 3 | 31 | 78626 |
| **B.catenulatum** | 100.0 | 43176 | 32372 | 16 | 216 | 17 | 78 | 23 | 189 |
| EU117603 | 100.0 | 40 | 33 | 39 | 23 | 12 | 16 | 246 | 73895 |
| **P.rettgeri** | 100.0 | 67274 | 54 | 17 | 258 | 6 | 159 | 27 | 67 |
| JF105442 | 100.0 | 29 | 79 | 62 | 22 | 4679 | 10 | 58538 | 190 |
| AY980174 | 100.0 | 29 | 55736 | 6 | 101 | 40 | 9 | 21 | 234 |
| HM310050 | 100.0 | 13 | 16 | 60 | 20 | 1524 | 4 | 51297 | 111 |
| **B.eggerthii** | 98.0 | 7815 | 41071 | 9 | 42 | 33 | 48 | 12 | 39 |
| HM278334 | 100.0 | 11 | 14 | 49 | 15 | 2249 | 5 | 46254 | 121 |
| **P.copri** | 97.0 | 4549 | 43031 | 3 | 23 | 53 | 210 | 16 | 14 |
| EU117683 | 100.0 | 22 | 16 | 20 | 17 | 11 | 12 | 242 | 45154 |
| KC001757 | 100.0 | 31 | 31 | 38537 | 51 | 11 | 8 | 89 | 441 |
| **R.faecis** | 100.0 | 6903 | 31029 | 9 | 132 | 85 | 31 | 23 | 36 |
| FJ504504 | 98.0 | 26 | 31359 | 5 | 107 | 52 | 6 | 14 | 50 |
| HQ242555 | 100.0 | 10 | 10 | 31274 | 16 | 13 | 4 | 17 | 111 |
| HQ671791 | 100.0 | 9 | 10 | 29779 | 49 | 7 | 3 | 119 | 68 |

|  |  |  |  |
| --- | --- | --- | --- |
| Genbank | Mock | Valid | Cross-talk |

To perform manual annotation, I examined each OTU in the table. If the lowest-abundance samples in a given OTU have much lower counts than the high-abundance samples, they are inferred to be probable cross-talk. In a mock sample, a high-abundance unexpected OTU, i.e. an OTU which does not match a species in the designed community, is probably a contaminant. A low-abundance unexpected mock count is probably cross-talk if it is also present in another sample. An alternative explanation is a low-abundance contaminant in the mock sample which is a valid OTU in the environmental samples by coincidence; this is a much less likely explanation. Another possible explanation is contamination which affects multiple samples, e.g. flow-cell residue from previous runs (Nelson *et al.*, 2014); this is also considered to be less likely than cross-talk. Under these assumptions, mock samples enable a more sensitive test for the presence of cross-talk. For example, if an unexpected mock OTU has two reads and some other sample has ten reads then the most likely explanation is cross-talk. The anomalously large cross-talk rate of 2/12 = 17% of the reads can be explained by fluctuations due to sampling effects when there are small total numbers of reads, which can result in high outlier values for some OTUs. In environmental samples, OTUs cannot be considered as expected or unexpected so abundances of two and ten in an OTU with twelve total reads is not a reliable indicator of cross-talk.

The UNCROSS algorithm described below uses simple heuristics to automate the manual procedure described above for annotating cross-talk. UNCROSS-Ref predicts cross-talk in mock samples using a reference database containing only expected sequences in the designed community. UNCROSS-Denovo predicts cross-talk in all samples considering read counts alone without using a database. These approaches are complementary. UNCROSS-Ref can identify unexpected OTUs by comparison with the database and is thus more sensitive to cross-talk in OTUs with low overall abundance, but cannot detect or correct cross-talk in environmental samples. UNCROSS-Denovo is less sensitive to cross-talk in OTUs with low overall read counts, but can detect cross-talk in environmental samples and can thus be used to detect and correct cross-talk in practice.

## UNCROSS algorithm

For a given OTU, let a *low* count be greater than zero and small enough to infer that most or all the reads for this sample are probably due to cross-talk. A *high* count is large enough to infer that most of the reads were correctly assigned to its sample. An *undermined* count is too large to be low and too small to be high (Fig. 2). UNCROSS uses simple heuristics to classify counts as low, undetermined or high. For a given OTU, variables are defined as follows.

> *S* is the number of samples.
>
> *N* is the total number of reads for all samples.
>
> *N_T_* is the number of reads which are assigned to the wrong sample.
>
> *M_T_* is the mean number of cross-talk reads per sample = *NT*/*S*. *R* is the *cross-talk rate* = *NT*/*N.*
>
> *n_H_* is the total number of reads in high counts, i.e. an estimate of the total number of reads in valid non-zero entries.
>
> *s_L_* is the number of samples with low counts.
>
> *n_L_* is the sum of low counts, i.e. an estimate of the total number of reads in counts which are non-zero due to cross-talk.
>
> *m_L_* is the largest low count.
>
> *m_avg_* is the mean low count = *nL*/*sL*.
>
> *m_max_* is the maximum number of reads assigned to a mock sample.
>
> *n_max_* is the maximum number of reads assigned to a non-mock sample. *mavg* is the mean number of reads assigned to a non-mock sample.
>
> *f_dn_* = *m_L_*/*N* is the *maximum cross-talk frequency* estimated by UNCROSS-Denovo.
>
> *f_ref_* = *m_max_*/*N* is the maximum cross-talk frequency estimated by UNCROSS-Ref.
>
> *r_ref_* = *S m_avg_*/*N* is the UNCROSS-Ref estimate of the cross-talk rate (*R*) (calculated only for OTUs where mock reads are predicted to be due to cross-talk).
>
> *r_dn_* is the UNCROSS-Denovo estimate of the cross-talk rate (*R*) (calculated only for OTUs where cross-talk is predicted).

Consider an OTU with 10 samples, three of which are mock. Suppose the counts are: mock = 200, 60, 10, other = 10000, 5000, 1000, 1000, 1000, 1000, 1000 for a total of *N* = 20270. The mock counts are low (probably cross-talk) and the rest are high (probably approximately correct). Then, *n_L_* = 270, *m_L_* = 200, *f_max_* = 200/20270 = 1%, *m_avg_* = 270/3 = 90 and *f_avg_* = *m_avg_*/*N* = 90/20270 = 0.45%. Some fraction of the reads assigned to samples with high counts will also be due to cross-talk, which can be estimated as follows. Assume *M_T_* is approximately *m_avg_* = 90. Then *N_T_* is approximately *S M_T_* = 900 misassigned reads and the estimated cross-talk rate *R* = *N_T_*/*N* = 900/20270 = 4.4%.

**Figure 1.**
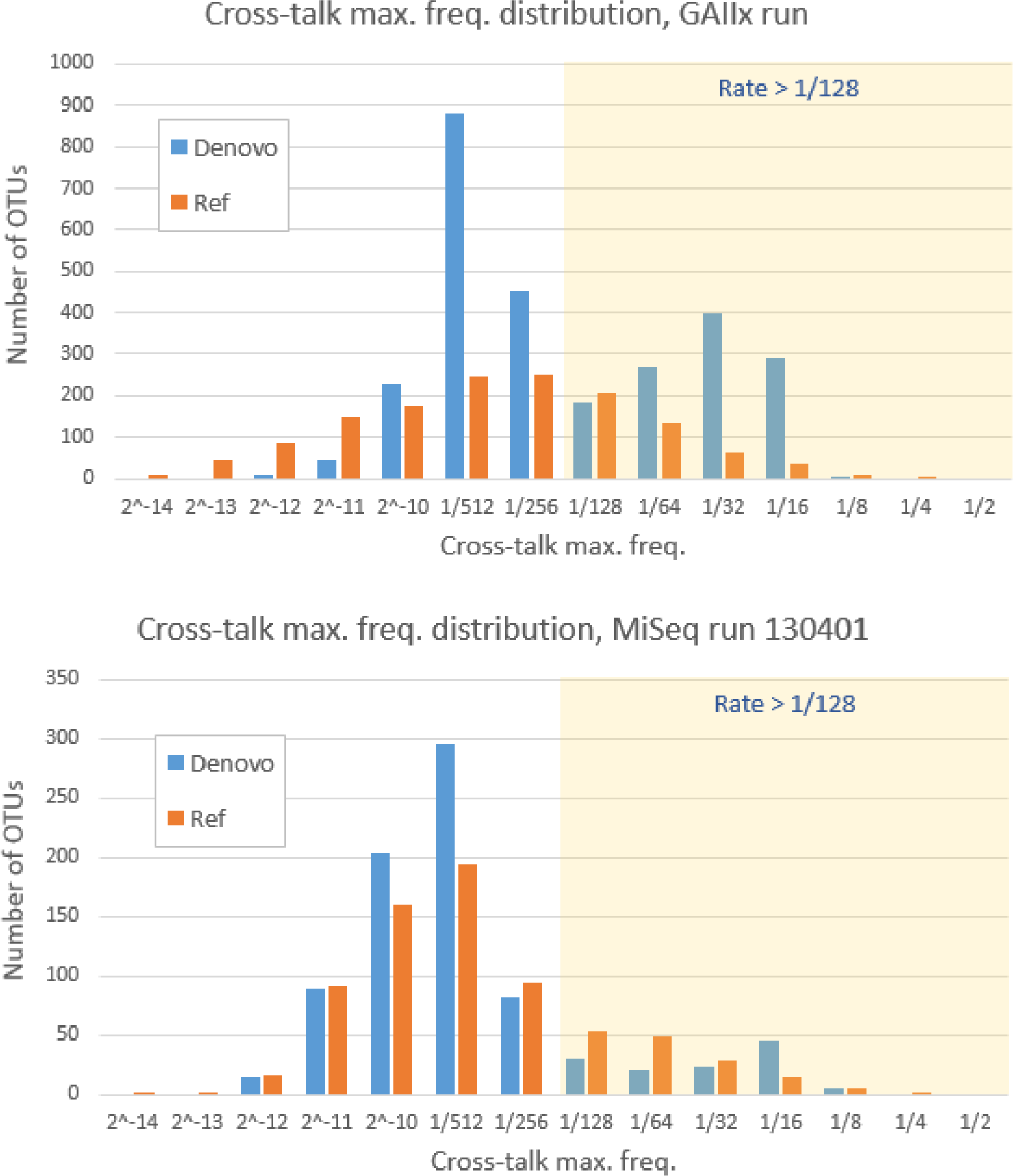
Cross-talk frequency distributions predicted by UNCROSS. Predicted maximum frequency for each OTU were assigned to bins with a minimum value shown in the horizontal axis, so the first bin contains OTUs with predicted rates from 2–14 to 2–13, the second bin from 2–13 to 2–12 and so on. Rates > 1/128 (i.e., more than ~1%) are highlighted. The maximum frequency is the largest OTU table entry predicted to be due to cross-talk divided by the total number of reads for the OTU. Since the number of cross-talk reads varies substantially between samples, the maximum frequency is the most relevant for setting a filter threshold.

**Figure 2.**
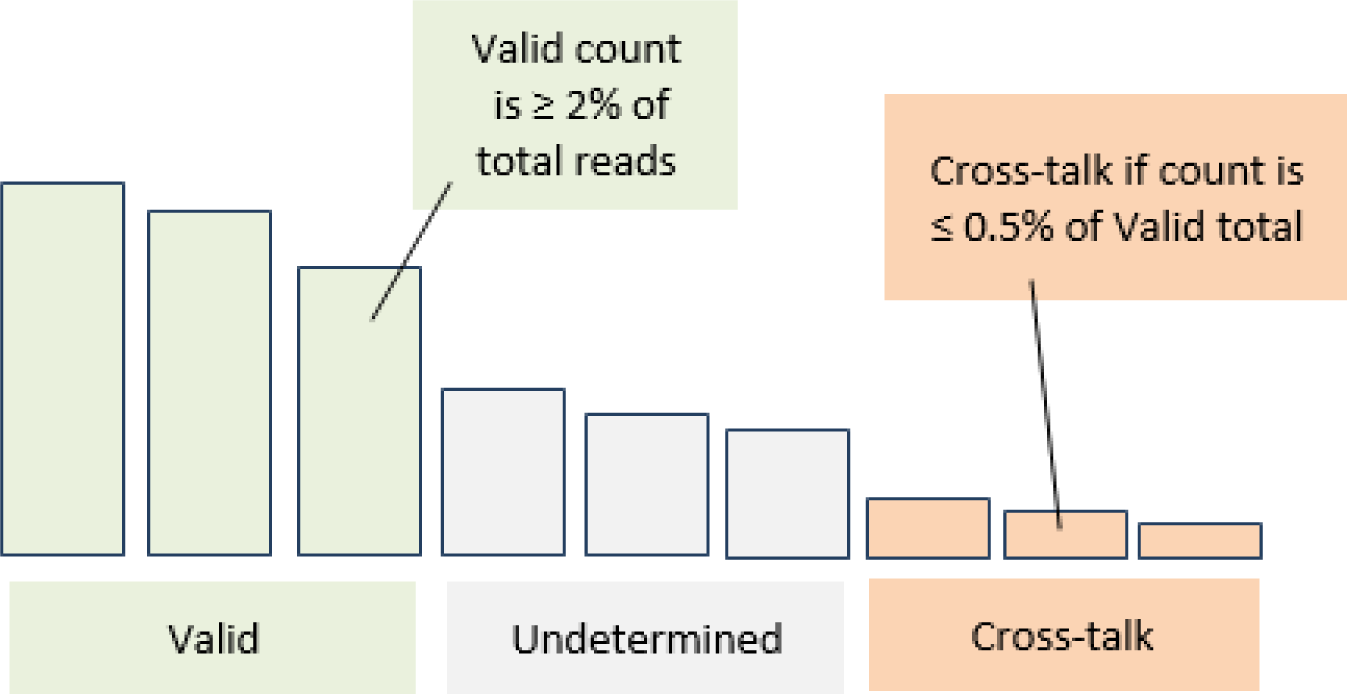
Schematic illustration of UNCROSS-Denovo algorithm. The OTU table entries for a given OTU are shown sorted by decreasing count (number of reads). If a count is at least 2% then it is classified as *valid*. If a count is = 0.5% of the total over *valid* counts, it is predicted to be due to cross-talk. Intermediate values are classified as *undetermined*.

### UNCROSS-Ref algorithm

The UNCROSS-Ref algorithm classifies an OTU as follows. If the total number of reads assigned to mock samples is zero, the OTU is not analyzed. If the sequence matches the reference database for the mock community, the OTU is classified as *designed*. Otherwise, the mock reads must be due to contamination or cross-talk, which is decided per the following pseudo-code.

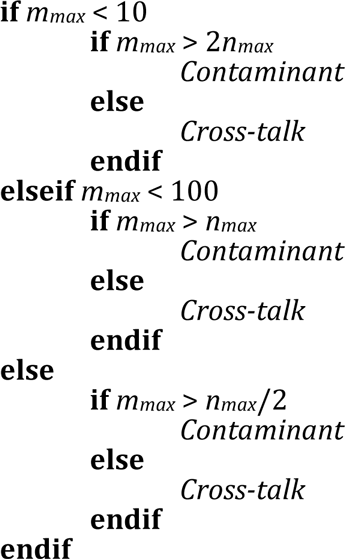

### UNCROSS-Denovo algorithm

The UNCROSS-Denovo algorithm classifies an OTU by considering the counts for each sample (Fig. 2). Non-zero counts are classified per the following rules.

A minimum value *v* for a valid count is calculated as follows: *if N* < 10 *then v*=5; *elseif N* < 100 *then v*=*N*/10; *else v*=*N*/50. Thus, for an OTU with at least 100 reads, a count of at least 2% of the reads is classified as *valid*. The sum of valid counts is *V*.

A maximum value *x* for a cross-talk count then is calculated as follows: *if V* < 10 *then x*=1; *elseif V* < 100 *then x*=*V*/10 + 1 *else x*=*V*/200. Counts which are neither valid nor cross-talk are classified as *undetermined*.

### UNCROSS-Denovo accuracy by comparison with UNCROSS-Ref

The accuracy of UCROSS-Denovo was assessed using UNCROSS-Ref as a gold standard by considering non-zero counts in the mock samples. Sensitivity was calculated by considering the subset of OTUs where the mock counts were predicted to be due to cross-talk by UNCROSS-Ref. Sensitivity is the fraction of these OTUs where all non-zero counts were also predicted to be cross-talk by UNCROSS-Denovo. The error rate was calculated as the fraction of all OTUs where the Ref and Denovo predictions disagreed on at least one mock sample. Results are shown in Table 3.

**Table 3.** Summary of results on twelve Illumina datasets. The first row is the GAIIx run from Caporaso *et al.*, the remaining eleven rows are MiSeq runs identified by run numbers from Kozich *et al.* Here, Ref means UNCROSS-Ref and Denovo means UNCROSS-Denovo. Columns are: *OTUs* number of OTUs, *Mock des.* number of OTUs matching designed mock strains, *Mock cont.* number of contaminant OTUs predicted by Ref, *Mock cross.* number of cross-talk OTUs predicted by Ref, *CX* number of non-zero counts (OTU table entries) predicted to be due to cross-talk (Denovo), *CU* number of undetermined non-zero counts (Denovo), *fdn* maximum cross-talk frequency (Denovo), *fref* maximum cross-talk frequency (Ref), *rref* estimated rate (Ref), *rdn* estimated rate (Denovo), *Acc*. accuracy of Denovo using Ref as a gold standard (fraction of mock predictions where Ref and Denovo agree).

| Run | OTUs | Mock<br>des. | Mock<br>cont. | Mock<br>cross. | $C_x$ | $C_U$ | $f_{ref}$ | $f_{dn}$ | $r_{ref}$ | $r_{dn}$ | Acc.. |
| --- | --- | --- | --- | --- | --- | --- | --- | --- | --- | --- | --- |
| GAllx | 7521 | 70 | 45 | 1407 | 32.6% | 23.9% | 0.39% | 0.44% | 2.62% | 10.41% | 75.1% |
| 121203 | 3591 | 18 | 1 | 202 | 7.5% | 27.2% | 0.26% | 0.23% | 1.78% | 2.07% | 86.1% |
| 121205 | 3630 | 18 | 2 | 249 | 8.3% | 26.2% | 0.26% | 0.24% | 1.75% | 2.21% | 84.7% |
| 121207 | 3560 | 18 | 1 | 182 | 7.3% | 26.9% | 0.29% | 0.24% | 2.02% | 2.09% | 86.3% |
| 130125 | 1134 | 19 | 1 | 28 | 4.2% | 29.7% | 0.67% | 0.26% | 2.09% | 2.36% | 114.3% |
| 130211 | 2650 | 19 | 1 | 164 | 6.1% | 25.7% | 0.33% | 0.27% | 1.76% | 2.45% | 79.9% |
| 130220 | 4156 | 18 | 1 | 374 | 9.5% | 27.6% | 0.24% | 0.22% | 1.42% | 2.27% | 82.6% |
| 130306 | 3504 | 18 | 1 | 464 | 11.3% | 27.0% | 0.34% | 0.26% | 1.25% | 2.52% | 72.8% |
| 130401 | 4378 | 18 | 1 | 708 | 14.7% | 26.3% | 0.27% | 0.25% | 1.14% | 2.13% | 78.4% |
| 130403 | 4455 | 19 | 1 | 696 | 14.4% | 26.2% | 0.26% | 0.25% | 1.12% | 2.01% | 77.3% |
| 130417 | 4339 | 19 | 1 | 524 | 12.4% | 26.3% | 0.24% | 0.24% | 1.12% | 2.02% | 78.6% |
| 130422 | 4252 | 19 | 1 | 454 | 11.3% | 26.6% | 0.26% | 0.26% | 1.46% | 2.19% | 78.6% |

## Discussion

In the data considered here, cross-talk is clearly identifiable in control samples of known composition (so-called mock communities). Unfortunately, mock samples are rarely included in practice. In fact, to the best of my knowledge, the runs analyzed here are the only public datasets where this type of analysis is possible. If cross-talk is present with frequencies comparable to those estimated here, diversity measures may be significantly degraded. Most OTUs assigned to the mock communities were spurious due to cross-talk, inflating OTU “richness” by an order of magnitude (Table 3). Alpha diversity metrics and estimators will be correspondingly inflated. Beta diversity measures will also be over-estimated if some samples have a long tail of shared but spurious OTUs which in fact are not present in those samples. These problems may be more serious when samples from distinctly different environments are sequenced in the same run. When samples from similar environments are compared, then cross-talk degrades the ability to make present / absent inferences for OTUs that have strongly varying abundance associated with certain metadata (e.g., before and after treatment with an antibiotic).

The UNCROSS algorithm uses simple heuristics that attempt to distinguish spurious OTU table entries due to cross-talk which should be zero from valid entries with low abundance. While UNCROSS works quite well on the datasets tested here, it was trained on the same data (because no other candidate training datasets are available, to the best of my knowledge) and it may be less accurate on other datasets. If there are no mock samples, and/or the number of samples is large, then automated *de novo* cross-talk detection may be more difficult or impossible, noting that cross-talk may have quite different rates and biases in other runs. If there are ~100, then the average OTU entry will have ~1% of the reads which is comparable to the maximum cross-talk frequency observed in the datasets tested here. This implies that cross-talk may be impossible to detect in most OTUs and that even present / absent inference for a given OTU in a given sample may be impossible in many or most cases. In conclusion, cross-talk is a well-documented but often neglected issue that should always be considered when analyzing multiplexed amplicon reads.

